# Meaning for reading pseudowords: errors reveal semantic influences on pseudoword reading after stroke

**DOI:** 10.64898/2026.05.13.724881

**Authors:** Ryan Staples, Elizabeth J. Anderson, Sara M. Dyslin, Alycia B. Laks, Rhonda B. Friedman, Andrew T. DeMarco, Peter E. Turkeltaub

## Abstract

Impaired reading, i.e., alexia, is common after left hemisphere stroke. The most common deficit in alexia is a difficulty reading aloud pronounceable novel words, also called pseudowords. While semantic and phonological processes both subserve reading real words, pseudoword reading deficits in alexia are typically ascribed to phonological deficits alone. Some theories, however, suggest that pseudoword reading relies in part on lexical-semantic knowledge, such that semantic deficits could also contribute to poor pseudoword reading in alexia. Leveraging a large sample of left-hemisphere stroke survivors, we examine the cognitive and neural substrates of pseudoword reading accuracy and two error types: lexicalization errors, where a pseudoword is incorrectly read as a real word, and nonword errors, where a pseudoword is read as an incorrect nonword.

76 left-hemisphere stroke survivors read 60 pseudowords aloud, and performed two pseudoword repetition tasks to assess phonological processing and two picture naming tasks to assess mappings between lexical semantics and phonology. Regression models assessed how pseudoword repetition and naming related to overall accuracy and rates of lexicalization and nonword errors in pseudoword reading. Voxel-based and connectome lesion-symptom mapping localized the neural territory responsible for these errors.

Pseudoword repetition and naming independently related to pseudoword reading accuracy. Pseudoword repetition but not naming deficits predicted higher rates of lexicalization errors, while naming but not pseudoword repetition deficits predicted higher rates of nonword errors. Greater nonword error rate also predicted smaller imageability effects in real word reading (*t*(71)=−3.2, *p*=0.002). Lexicalization errors were associated with lesions to and disconnections of the left putamen and basal ganglia. Nonword errors were associated with lesions to the superior and middle temporal gyri, as well as broad temporo-parietal disconnections, overlapping with previous lesion-mapping results implicating these regions in semantic contributions to word reading. These results suggest that lexicalization errors result from impaired planning and execution of novel motor plans, causing a reliance on the well-learned motor plans associated with lexical items. In contrast, greater rates of nonword errors, relative to lexicalization errors, occur when semantic contributions to reading are impaired.

Overall, these findings demonstrate that semantic processes are involved in reading pseudowords, at least in stroke alexia. These findings support connectionist accounts of reading in which damage in the direct orthography to phonology route for reading leads to greater reliance on semantic representations, even for pseudowords, suggesting a reinterpretation of pseudoword reading as a pure measure of phonological deficits in reading.

## Introduction

Two-thirds of left hemisphere stroke survivors (LHSS) with aphasia also have alexia, an acquired disorder of reading.^1^ Literacy is essential in contemporary life, and persons with alexia report that impaired reading causes severe negative impact on well-being.^2^ A better understanding of the neurocognitive basis of alexia is thus critical to informing interventions to restore reading after stroke.

Behavioral, computational, and neural evidence have converged on a two-stream neurocognitive model of oral reading. Early neuropsychological studies demonstrated that the reading of irregular words (words that do not follow the standard spelling-to-sound correspondences of English) and pseudowords (pronounceable, graphotactically correct novel letter strings that are not meaningful) can be separately impaired.^3–5^ These findings informed the development of cognitive models positing that letter strings are read by interactive computation between a sublexical print-to-sound decoding stream, subserved by phonological processing^6^, and a whole-word stream^7,8^, driven by lexical-semantic processes.^9^ Based on these cognitive models, difficulty reading pseudowords relative to real words is diagnostic of a phonological reading deficit resulting from damage or dysfunction in the sublexical stream. In the syndromic classification scheme of alexias, this type of deficit is referred to as phonological alexia, which is the most common type of post-stroke alexia.^1^

The dual stream cognitive architecture of reading maps onto an observed neurological distinction in speech processing: a ventral stream, mapping between wordforms and meaning, and dorsal stream, mapping between wordforms and articulation.^10,11^ Dorsal stream cortical regions include the superior temporal gyrus (STG), the supramarginal gyrus (SMG), and the premotor cortex (PMC).^12,13^ The left STG is thought to store phonological representations, ranging from low level phonemic representations to larger auditory wordform representations^14–17^. The SMG is likely involved in auditory working memory and in articulatory sequencing, critical for online assembly of pseudoword motor plans.^18^ Finally, the motor plans for speech are thought to be stored in the posterior inferior frontal gyrus (IFG) and premotor cortex.^13,19,20^ Subcortically, the basal ganglia play a critical role in motor action selection and control.^21,22^ Evidence suggests a caudate-gated planning loop, responsible for selecting an upcoming syllable or phoneme, and a putamen-gated motor loop that ‘releases’ planned speech programs to the articulators.^21^ Damage to dorsal stream regions causes a variety of speech disorders. Lesions to the SMG cause conduction aphasia, where repetition is impaired but auditory comprehension is relatively spared.^23,24^ Lesions to the STG impair acoustic-phonetic perception and can cause pure word deafness.^25^ Posterior IFG and PMC damage cause apraxia of speech, a disorder of motor planning.^26–28^ Finally, subcortical damage is most associated motor speech impairment.^29–31^ Critically, lesions to dorsal stream regions disproportionately impair sublexical processing, leading to worse pseudoword reading relative to real word reading.^32,33^ Further, damage to different dorsal stream processors results in subtly different patterns of phonological reading deficits within the broader diagnosis of phonological alexia.^32,33^

Neurocognitive models of oral reading suggest that pseudowords primarily rely on the dorsal stream^7,34,35^: their pronunciations are assembled from sublexical orthography-to-phonology decoding, and the computed wordform is passed to articulatory cortex. However, computational ‘triangle’ connectionist models of reading learn sublexical associations from phonology and orthography to semantics, and thus activate semantics when processing pseudowords.^36^ Does the lexical-semantic route for reading then contribute to pseudoword reading? The pseudohomophone effect, wherein pseudowords that can be pronounced like real words (e.g. *brane*, /breLn/) are read more quickly than pseudowords that cannot^37,38^, provides evidence that lexical phonology, and perhaps lexical semantics, are activated by meaningless letter strings.^37^ However, persons with severe semantic impairment due to semantic-variant primary progressive aphasia (svPPA) often have preserved pseudoword reading.^9^ This suggests that if semantic processing contributes to pseudoword reading, it is not necessary, at least in the absence of phonological deficits. Likewise, damage to ventral stream regions, including the left fusiform (FG), middle temporal (MTG), and angular gyri (AG), as well as the ventral anterior temporal lobe (ATL),^11,12,39^ has been associated with lexical and semantic but not pseudoword reading deficits. In stroke alexia, where phonological processing is often impaired, the recruitment of intact lexical-semantic processes may be upregulated, even for the reading of pseudowords. We recently showed that lesions to the posterior superior temporal sulcus (pSTS) were associated with both impaired reading of high imageability words and picture naming, suggesting an essential role in mediating the semantic-phonological mapping critical for reading.^40^ The pSTS is thus an excellent candidate for mediating such an effect in pseudoword reading.

While most studies of pseudoword reading deficits focus on accuracy, the types of errors made in pseudoword reading may provide a finer-grained measure of cognitive influences on reading than accuracy alone. We organize errors into two broad categories. Implausible nonword errors (NE) are those in which a pseudoword is read as another pseudoword that is not a plausible pronunciation given the spelling (‘clais. -> /klest/). Lexicalization errors (LE), in contrast, are those in which a pseudoword is read aloud as a real word that is implausible given the spelling (‘yoog’ -> yogurt).

Lexicalizations comprise a large percentage of pseudoword reading errors in LHSS^5^, a population that is usually phonologically impaired due to the proximity of the middle cerebral artery to dorsal stream regions.^1^ One study of LHSS found that the rate of LE and a composite score of semantic performance were positively correlated, suggesting that semantic processes contribute to pseudoword reading.^41^ However, that study utilized a correlational analysis that did not account for both phonological and semantic impairment simultaneously, which is critical given that phonological impairments result in poor pseudoword reading.^33,42^ A computational modeling study that examined reorganization of reading pathways after lesions suggested a mechanism for LE: phonological lesions to the model reweighted processing during plasticity-related recovery to be more semantically-mediated in nature. LE rates rose only after this reorganization occurred.^43^ While not specific to pseudowords, direct support for semantic compensation in oral real word reading is provided in a pair of fMRI studies finding selectively increased activation in the left hemisphere semantic network on accurate oral reading trials, relative to inaccurate trials, in phonologically-impaired stroke surviviors.^44,45^ These studies converge to suggest that when phonology is impaired, readers rely on the semantically-mediated pathway – if it is intact. LE are thus likely to occur in stroke survivors with impaired phonology but intact semantics, due to a reliance on lexical reading.

The basis of NE in pseudoword reading is less clear. Impaired sublexical processing is a likely source of these errors, and previous lesion studies support that SMG lesions impair pseudoword reading accuracy.^32,33^ However, no direct evidence speaks to the cognitive or neural drivers of NE relative to LE. We suggest that when both phonology and semantics are impaired, readers must rely on a damaged sublexical assembly process when reading pseudowords, producing a relatively greater rate of NE.

If rates of NE, relative to LE, reflect damage to underlying lexical-semantic processes, then they should also relate to semantic effects in real word reading. A common index of semantic processing in reading is the imageability effect: words that are imageable (they evoke a clear mental picture, e.g. ‘hammer’) are read more quickly than those that are not (‘justice’).^46,47^ We previously found that impaired reading of high relative to low imageability words was related to picture naming impairment.^40^ Here, we extend this finding to ask if NE errors in pseudoword reading predict high imageability word reading accuracy.

No prior studies have examined the neural basis of pseudoword reading error types, or simultaneously assessed how semantics and phonology contribute to pseudoword error type production. We characterize the rates of LE and NE during pseudoword reading in large dataset of LHSS, and examine if phonological or semantic processes impact the production of these error types. Building on our previous work, we then test if PE and LE rates predict the reading of high imageability words.^40^ Finally, we use support-vector regression voxel-based lesion-symptom mapping (SVR-VLSM) and connectome lesion-symptom mapping (SVR-CLSM) to identify the local and network neural correlates of error type in pseudoword reading. We predicted that both phonological and semantic impairments relate to pseudoword reading deficits, with relative impairments reflected in error patterns. Impaired phonological processing and damage to the dorsal stream would cause greater rates of LE production, due to a reliance on lexical reading processes. Impaired semantic processes in the ventral stream would result in greater relative rates of NE production, due to a reliance on sublexical assembly.

## Materials and methods

### Participants

Participants were 76 LHSS and 62 age- and education-matched neurotypical controls (Table 1). All participants were recruited as part of an ongoing study on individual differences in aphasia after left-hemisphere stroke (clinicaltrials.gov NCT04991519). Participants were native English speakers and had adequate vision and hearing with correction from lenses or hearing aids to complete auditory and visual tasks. Controls were excluded if they scored below an age, ethnicity, and race^48^ adjusted cutoff on the Montreal Cognitive Assessment.^49^ Exclusion criteria for both the stroke and control cohorts were: history of significant neurological disease, head injury causing loss of consciousness, learning disorder requiring educational intervention, psychiatric disorder requiring hospitalization, or ongoing use of psychiatric medications other than common antidepressants. All participants in the stroke cohort were in the chronic stage of recovery (>six months post-stroke) at the time of enrollment. To ensure that sufficient verbal output was present in all participants, we removed 12 stroke survivors who had severe apraxia of speech per the Apraxia of Speech Ratings Scale 3.0 (see below).^50,51^ Additionally, six stroke survivors were removed for not completing the pseudoword reading task. The final sample consisted of all the controls and stroke survivors from within the recruitment period who fulfilled these criteria. Four controls were excluded in order to achieve demographic matching between groups. Written informed consent was obtained from all participants as required by the Declaration of Helsinki. The study protocol was approved by the Georgetown University Institutional Review Board. Controls were used solely in deriving normative connectomes.

**Table 1.**
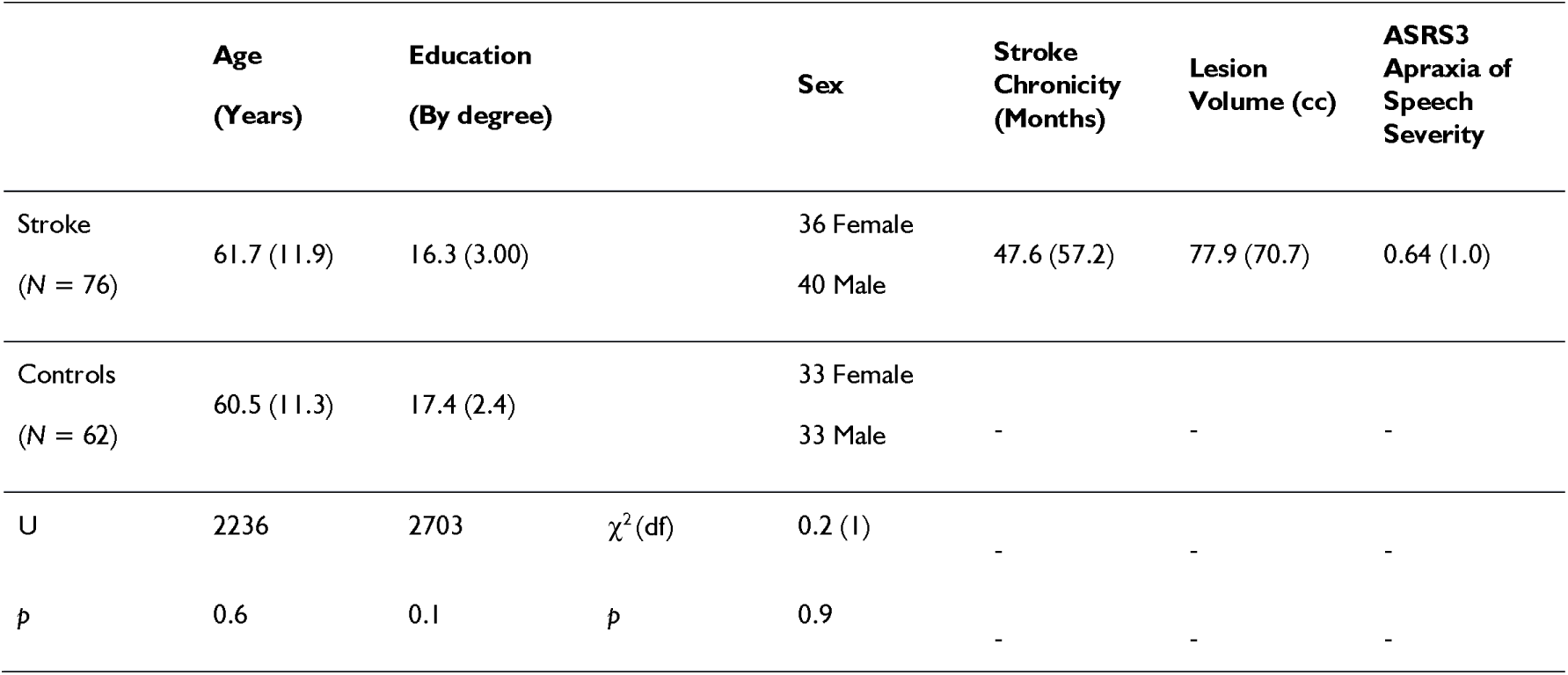
Participant demographics and clinical data.

|  | Age<br>(Years) | Education<br>(By degree) |  | Sex | Stroke<br>Chronicity<br>(Months) | Lesion<br>Volume (cc) | ASRS3<br>Apraxia of<br>Speech<br>Severity |
| --- | --- | --- | --- | --- | --- | --- | --- |
| Stroke<br>(N = 76) | 61.7 (11.9) | 16.3 (3.00) |  | 36 Female<br>40 Male | 47.6 (57.2) | 77.9 (70.7) | 0.64 (1.0) |
| Controls<br>(N = 62) | 60.5 (11.3) | 17.4 (2.4) |  | 33 Female<br>33 Male | - | - | - |
| U | 2236 | 2703 | $\chi^2$ (df) | 0.2 (1) | - | - | - |
| p | 0.6 | 0.1 | p | 0.9 | - | - | - |

### Behavioral Testing

#### Pseudoword Reading

Participants read aloud a list of 60 pseudowords. All stimuli were monosyllabic and 3-6 letters in length. Twenty pseudowords had only one plausible pronunciation given orthographic bodies in English, 20 had two or more plausible pronunciations, and 20 pseudowords had orthographic bodies with no comparable mapping. Participants had 10 seconds to read aloud each pseudoword, after which the trial timed out. The first complete attempt containing both a consonant and vowel was scored. See Dyslin et al.^32^ for further details of the stimuli and presentation. Any pronunciation that could plausibly be produced by a spelling-to-sound correspondence present in English was scored correct. Thus, both the LE and NE we analyze are exclusively incorrect pronunciations given the spelling. Unintelligible responses, as well as other errors such as omissions and stereotypy responses, were discarded. We use error rates across all analyses of error types, calculated as the number of trials in which an error of a given type was elicited, divided by 60 (the number of pseudowords read).

#### Real Word Reading

We assessed semantic influences on reading by examining differences between high and low imageability word reading. Participants read aloud 200 English words, categorically divided into high/low imageability, high/low frequency, and regular/irregular lists. There were 100 words in each list. High and low imageability words were matched on letter length, frequency, regularity, and articulatory complexity. Participants were told to read each word aloud as quickly and accurately as possible within a 10 second time limit. The first attempt containing a vowel and consonant (CV or VC) was scored, or an isolated vowel if the target word was a CV or VC. Further details of the corpus and test administration can be found in Staples et al.^40^

#### Pseudoword Repetition

To assess phonological processing ability, two pseudoword repetition tasks were used to probe phonological processes in the LHSS. In the first, participants were presented with 30 pseudowords auditorily and were required to repeat them aloud. Items varied on the number of syllables, with 10 items at each length: one, three, and five syllables. If they could not repeat the word within 10 seconds, they timed out and the next trial began. Trials were scored as correct if the elicited pronunciation matched the cue exactly.^52^ The second pseudoword repetition task had the same task structure, but with 60 trials.^53^ Again, items varied on the number of syllables, with 20 items at each length: one, two, and three syllables.

#### Picture Naming

Previously, we found that semantic reading deficits in alexia, as indexed by reduced imageability effects, are related to deficits in semantic-to-phonological mapping, as indexed by picture naming.^40^ As in that paper, we assess the intactness of semantic-to-phonology mappings by averaging accuracy on two different picture naming tasks: the 60-item Philadelphia Naming Test^54^ and an in-house 60-item picture naming test.^55^ In both tasks, participants were instructed to use one word to name aloud each picture as quickly and accurately as possible, within a 20-second limit. Accuracy was scored on the first complete attempt.

#### Apraxia of Speech Assessment

The Apraxia of Speech Ratings Scale 3.0 (ASRS3) was used to assess stroke survivors for motor speech impairment. Stroke survivors were rated on severity using a scale of 0 (indicating “not present) to 4 (“nearly always evident”) ^50,51^. A certified speech-language pathologist performed ratings via video review. Stroke survivors with motor speech impairments were given leniency on their oral responses for all tasks: credit was given if their response included a prolongation, segmentation, or distortion that did not cross phoneme boundaries. As noted above, individuals with severe apraxia were excluded from the study because the severity of apraxia precluded reliable error coding.

### Neuroimaging

#### MRI Acquisition and Lesion Tracing

The following sequences were acquired with Georgetown’s 3T Siemens MAGNETOM Prisma scanner using a 20-channel head coil: a T1-weighted magnetization prepared rapid gradient echo (MPRAGE) sequence (1mm^3^ voxels), a fluid-attenuated inversion recovery (FLAIR) sequence (1mm^3^ voxels), and a high angular resolution diffusion imaging (HARDI) sequence (81 directions at *b*=3000, 40 at *b*=1200, 7 at *b*=0; 2 mm^3^ voxels). For LHSS, author P.E.T., a board-certified neurologist, manually traced stroke lesions on the native-space MPRAGE and FLAIR images via ITK-SNAP^56^; http://www.itksnap.org/). MPRAGEs and lesion tracings were warped to the Clinical Toolbox Older Adult Template^57^ using Advanced Normalization Tools ^58^ as described in Dickens et al.^59^

#### Structural Connectome Pipeline

MRtrix 3.0^60^ software was used to preprocess the HARDI data, estimate white matter pathways, and create a structural connectome for each subject. Multi-shell, multi-tissue constrained spherical deconvolution^61^ was applied to calculate the voxelwise fiber orientation distributions from the preprocessed HARDI data. Probabilistic anatomically constrained tractography^62^ on the white matter fiber orientation distributions traced 15 million streamlines in native space (algorithm=iFOD2, step=1, min/max length=10/300, angle=45, backtracking allowed, dynamic seeding, streamlines cropped at grey matter-white matter interface). The Spherical Deconvolution Informed Filtering of Tractograms-2 algorithm^63^ was applied to derive cross-sectional multipliers for each streamline to adjust streamline densities to be proportional to the underlying white matter fiber densities. Subject-wise white matter connectivity matrices were created by assigning streamlines to the 246 brain parcels of the Brainnetome atlas.^64^ Each edge (connection) of a connectome represents the apparent fiber density of the white matter connecting the two brain parcels. For the purpose of SVR-CLSM, each LHSS’s connectome was binarized by comparing stroke and control connectomes to label each connection as lesioned (0) or intact (1). Specifically, a connection in an LHSS’s connectome was deemed lesioned if the apparent fiber density was below the first percentile of the control values for that specific connection. Further details of the connectome pipeline can be found in Dickens et al.^33^

### Statistical Analyses

#### Assessing semantic and phonological processing

To establish if stroke survivors were more semantically or phonologically impaired, a repeated-measures ANCOVA predicted accuracy from task (pseudoword repetition vs. picture naming), covarying for age, education, log-transformed chronicity in months, ASRS AOS Severity, and lesion volume.

#### Relating pseudoword reading accuracy and error types to semantic and phonological processing

To assess if both semantic and phonological processing contribute to pseudoword reading, a linear regression estimated participant-wise pseudoword reading accuracy from pseudoword repetition and naming accuracy, with age, education, log-transformed chronicity in months, ASRS AOS Severity, and lesion volume as additional predictors. Lesion volume and AOS severity were included to examine effects of overall impairment and articulatory impairment. Chronicity was included to test if pseudoword reading ability relates to post-stroke recovery over time.

Two separate linear regressions models assessed how semantic and phonological processes contribute to 1) LE and 2) NE. The first model predicted LE rate as a function of accuracy on pseudoword repetition and naming accuracy, with NE rate and the covariates listed above as additional predictors. The second model predicted pseudoword rate from pseudoword repetition and naming accuracy, with LE rate and the same covariates as additional predictors. Including the opposite error type as a covariate (e.g., including NE rate as a covariate for LE) controls for overall performance as well as the moderate correlation between error types.

#### Relating pseudoword reading errors to real word processing

To relate pseudoword error rates to semantic influences on real word reading, a linear regression model predicted high imageability word reading from PE and LE rates, accuracy on low imageability word reading, and the demographic variables listed above. Including low imageability words as a predictor isolates the imageability effect.

#### Voxel-based lesion-symptom mapping

To localize lesions that impair pseudoword reading, we conducted an SVR-VLSM analysis of pseudoword reading accuracy, covarying for age, education, ASRS3 AOS Severity, and lesion volume.

We then localized lesions that are specifically related to LE and NE production. Two SVR-VLSM analyses identified where damaged tissue related to 1) LE production rates and 2) NE production rates.^65^ The LE rate analysis covaried for NE rate, and vice versa, controlling for overall performance and ensuring the analyses identify tissue that is uniquely associated with each error type. Age, education, ASRS3 AOS Severity, and lesion volume were regressed out of both the behavior and lesion data prior to SVR.

Only voxels lesioned in at least 10% of the participants (*n*=8) were included in the analyses. Permutation tests (5000 permutations), shuffling behavioral scores among participants, were used to identify significant results at a voxelwise *p*<0.005 and family-wise error rate of *p*<0.05.^66^ All analyses were performed using the *svrlsmgui* function from the svrlsmgui toolbox (https://github.com/atdemarco/svrlsmgui)^67^ for MATLAB.

#### Connectome lesion-symptom mapping

We conducted two SVR-CLSM analyses to identify disconnections that caused increased rates of NE and LE using an in-house custom MATLAB script. These analyses were analogous to the two SVR-VLSM analyses, with the same covariates. We also estimated an SVR-CLSM analysis of pseudoword reading accuracy, exactly analogous to the SVR-VLSM analysis described above. Only connections present in 100% of the control participants were included, reducing Type 1 error. Only connections within the left hemisphere and between the left and right hemisphere were assessed, because all participants had left-lateralized strokes. Only connections lesioned within at least 10% of participants (*n*=8) were included in the analysis. Edge-wise significance was calculated via 10000 permutations, with continuous family-wise error rate control using v=10 and family-wise error rate=0.05, one-tailed (negative).^68,69^ As in the SVR-VLSM analysis, disconnections that are specific to an error type and do not overlap with the accuracy result should identify white matter connections that are unique to producing error types.

#### Connectome lesion-symptom mapping

We conducted two SVR-CLSM analyses to identify disconnections that caused increased rates of NE and LE using an in-house custom MATLAB script. These analyses were analogous to the two SVR-VLSM analyses, with the same covariates. We also estimated an SVR-CLSM analysis of pseudoword reading accuracy, exactly analogous to the SVR-VLSM analysis described above. Only connections present in 100% of the control participants were included, reducing Type 1 error. Only connections within the left hemisphere and between the left and right hemisphere were assessed, because all participants had left-lateralized strokes. Only connections lesioned within at least 10% of participants (*n*=8) were included in the analysis. Edge-wise significance was calculated via 10000 permutations, with continuous family-wise error rate control using v=10 and family-wise error rate=0.05, one-tailed (negative).^68,69^ As in the SVR-VLSM analysis, disconnections that are specific to an error type and do not overlap with the accuracy result should identify white matter connections that are unique to producing error types.

## Results

### Rates of pseudoword errors types

We found that most stroke survivors (69/76) made both NE and LE (Fig. 1A). Correlations showed that LE and NE rates were positively related (*r*=0.34, *t*(74)=3.1, *p*=0.003). However, there remained wide individual variation in error type rates (Fig. 1F).

**Fig. 1.**
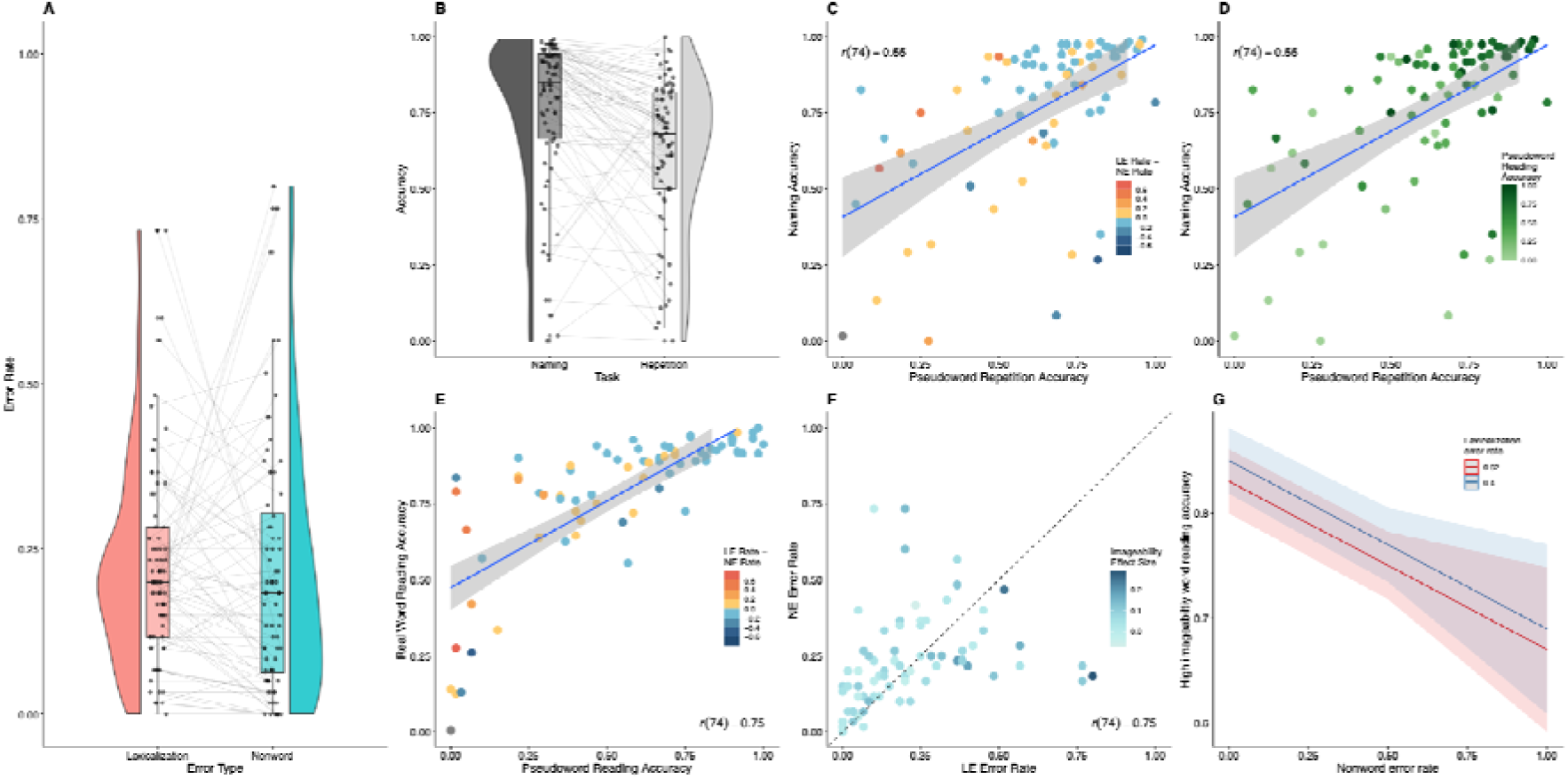
Plots showing behavioral performance in stroke survivors. **(A)** Rates of and lexicalization and nonword error rates in the pseudoword reading task. **(B)** Performance on pseudoword repetition and naming. Lines connect data points for individuals. Horizontal jitter added to the scatter plots for error rates and task performance disambiguate overlapping data points. **(C)** Relationship between performance on picture naming, pseudoword repetition, and rates of error types. Points are colored by a subtraction of lexicalization error rate – nonword error rate. Warm colors indicate participants who make more lexicalizations than nonword errors. Cool colors indicate participants who make more nonword than lexicalization errors. **(D)** Relationship between picture naming, pseudoword repetition, and pseudoword reading accuracy. Darker color indicates better pseudoword reading accuracy. **(E)** Relationship between word and pseudoword reading. Points are colored by a subtraction of lexicalization error rate – nonword error rate. Hot colors indicate participants who make more lexicalizations than nonword errors. Cool colors indicate participants who make more nonword than lexicalization errors. The plot suggests that individuals with poor word reading tend also have poor pseudoword reading. Larger relative deficits of pseudoword reading were associated with more lexicalization errors, while larger relative word reading deficits are associated with more nonword errors. **(F)** Relationship between nonword and lexicalization error rates. Points are colored by the size of the imageability effect (high imageability word reading accuracy – low imageability word reading accuracy). Darker colors indicate larger imageability effects. The plot suggests that individuals with large imageability effects tend to make more lexicalization errors while those with lower imageability effects make more nonword errors. **(G)** Results of a regression model predicting high imageability word reading accuracy from lexicalization and pseudoword error rates, controlling for low imageability word reading accuracy, age, education, lesion volume, log-transformed chronicity in months, and ASRS3 Apraxia of Speech Severity. Lines indicate +/-1 SD lexicalization error rate. Nonword errors, but nor lexicalization errors, predicted high imageability word reading.

**Table 2.**
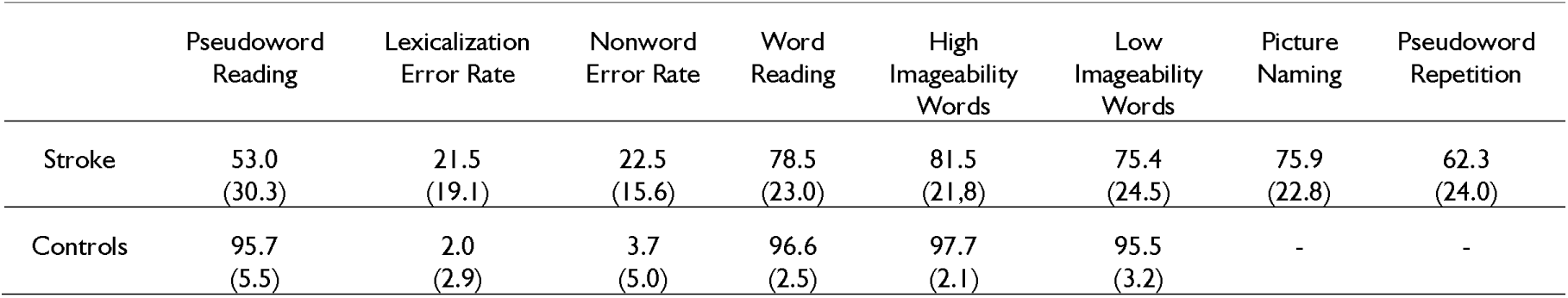
Performance on behavioral tasks.

|  | Pseudoword Reading | Lexicalization Error Rate | Nonword Error Rate | Word Reading | High Imageability Words | Low Imageability Words | Picture Naming | Pseudoword Repetition |
| --- | --- | --- | --- | --- | --- | --- | --- | --- |
| Stroke | 53.0<br>(30.3) | 21.5<br>(19.1) | 22.5<br>(15.6) | 78.5<br>(23.0) | 81.5<br>(21.8) | 75.4<br>(24.5) | 75.9<br>(22.8) | 62.3<br>(24.0) |
| Controls | 95.7<br>(5.5) | 2.0<br>(2.9) | 3.7<br>(5.0) | 96.6<br>(2.5) | 97.7<br>(2.1) | 95.5<br>(3.2) | - | - |

### Characterizing phonological and semantic processing in stroke survivors

Performance on the naming and pseudoword repetition tasks were correlated (*r*=0.55, *t*(74)=5.7, *p*<.001), but there was large variability (Fig. 1C,D). A repeated-measures ANCOVA found that, as a group, stroke survivors were more impaired in pseudoword repetition than naming (F(1,74)=4.71, *p*<.033), indicating worse phonological than semantic impairment (Fig 1B; Supplementary Table 1). Lesion volume (F(1,74)=15.56, *p*<.001), ASRS3 AOS Severity (F(1,74)=18.58, *p*<.001), and chronicity (F(1,74)=5.84, *p*=.018) were also predictive of task performance.

### Identifying the cognitive drivers of pseudoword reading and error rates

A linear regression model found that better pseudoword reading accuracy was predicted both by better pseudoword repetition (β=0.25, *t*(67)=2.07, *p*=.042; Table 3) and naming scores (β=0.51, *t*(67)=4.32, *p*<.001; Table 3; Fig. 1D). Larger lesion volume (β=−0.00, *t*(67)=−2.80, *p*=.007) and longer chronicity (β=−0.07, *t*(67)=−3.05, *p*=.003) both predicted worse pseudoword reading accuracy (overall regression: adjusted *R*^2^=0.61, *F*(7,68)=17.44, *p*<.001; Table 3).

**Table 3.**
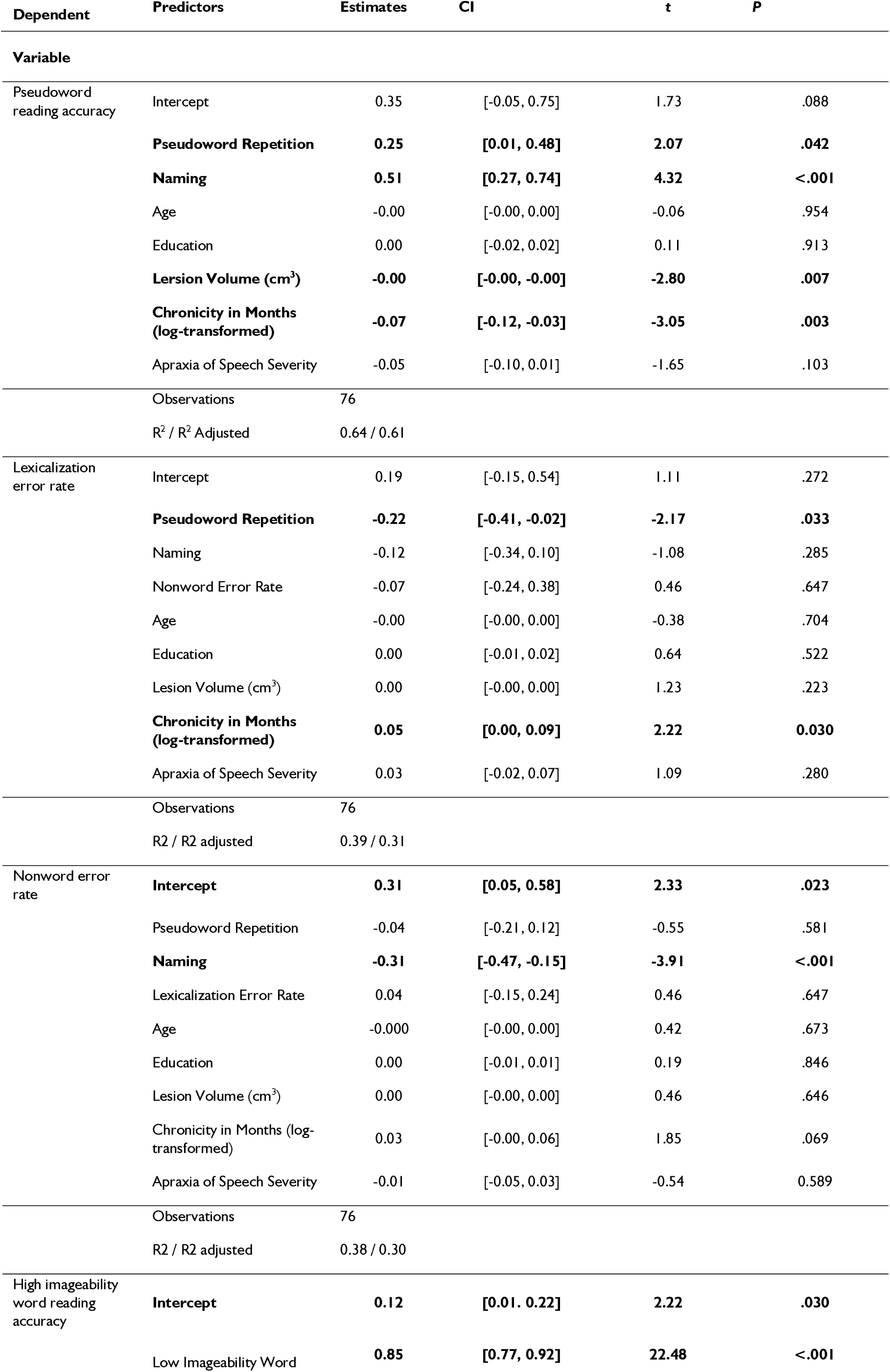
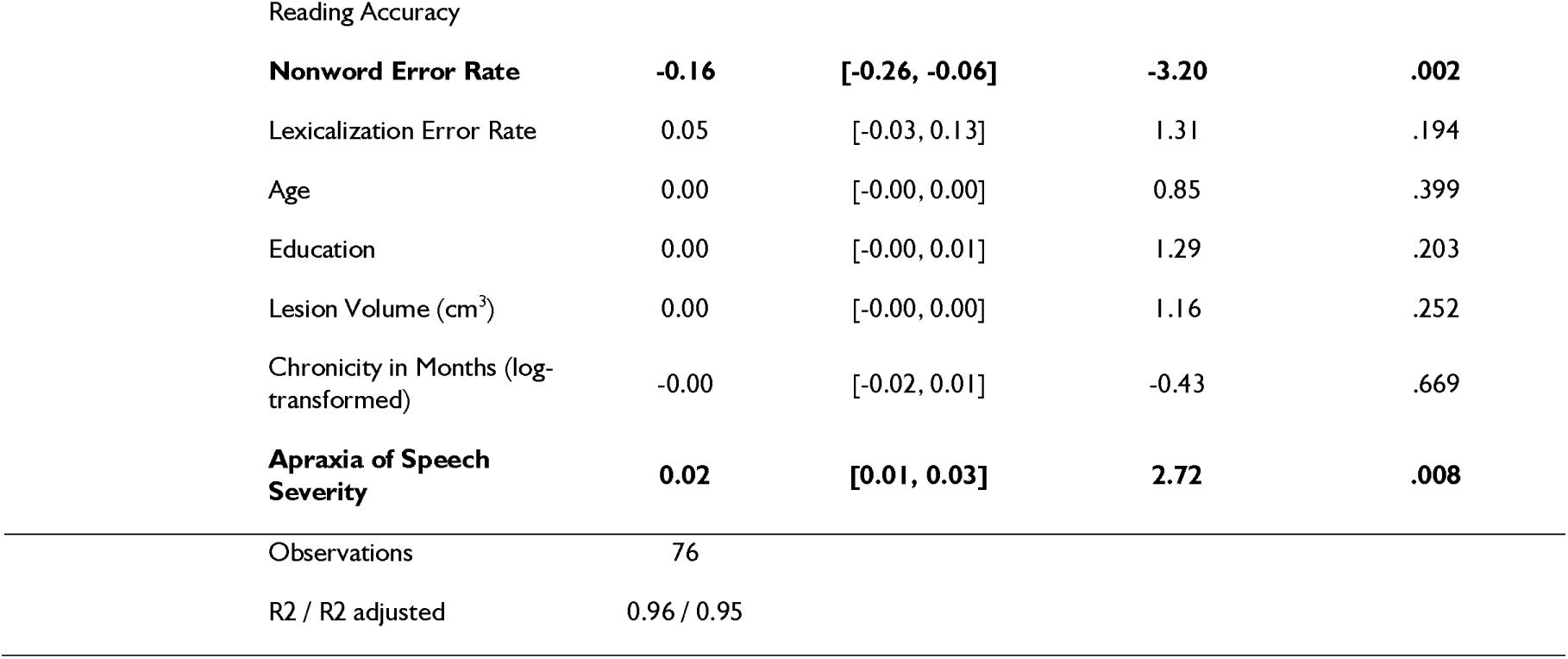
Results of regression models.

Two linear regression models tested if LE and NE production were predicted by phonological and semantic processing. LE rate was predicted by pseudoword repetition (β=−0.22, *t*(67)=−2.17, *p*=.033; Fig. 1C) but not naming, indicating that worse phonological processing contributed to producing more LE (overall regression: adjusted *R*^2^=0.31, *F*(8,67)=5.24, *p*<.001; Table 3). We also found that longer chronicity was associated with larger rates of LE (β=0.05, *t*(67)=2.22, *p*=.030). Greater NE rate was predicted by worse naming (β=−0.31, *t*(67)=−3.91, *p*<.001; Fig. 1C) but not pseudoword repetition (overall regression: adjusted *R*^2^=0.30, *F*(8,67)=5.05, *p*<.001; Table 3). There was also a trend towards longer chronicity predicting larger NE rates (β=0.03, *t*(67)=−1.85, *p*=.069).

### Relating pseudoword error rates to oral reading

Pseudoword and real word reading were strongly correlated (*r*=0.75, *t*(74)=9.7, *p*<.001; Fig. 4E). Visual inspection shows that larger deficits of pseudoword reading are associated with greater rates of LE, indicating a reliance on lexical-semantic reading when phonology is impaired. More severe word reading deficits are instead associated with a larger NE rate, suggesting a reliance on noisy sublexical decoding (Fig 4E).

The results above suggest that semantic processing deficits plays a role in pseudoword reading, particularly in the production of NE. If so, then reduced reliance on semantics for word reading, as evidenced by loss of the advantage for reading high imageability words relative to low imageability words, should relate to production of NE in pseudoword reading (Fig. 4F). In a regression testing this prediction, greater NE error rates but not LE rates predicted worse high imageability word reading (β=−0.16, *t*(65)=−3.20, *p*=.002; Fig. 4E-G). High imageability word reading was also predicted by better low imageability word reading (β=0.85, *t*(65)=22.48, *p*<.001) and greater AOS severity (β=0.02, *t*(65)=2.72, *p*=.008).

### Identifying the neural loci of pseudoword errors

Lesion coverage was robust throughout the perisylvian region, with maximal lesion overlap located in the insula and precentral gyrus (Max. overlap=38; Fig. 2A). SVR-VSLM identified a cluster associated with reduced pseudoword reading accuracy, located in the posterior superior temporal gyrus and supramarginal gyrus (MNI centroid: −51.7, −36.2, 17.4; Fig. 2B; Table 4). Examination of the sub-threshold Z maps suggested that beyond this focal area, damage to neighboring motor, parietal, fusiform, and frontal cortex were weakly associated with reduced pseudoword reading accuracy (Supplementary Fig. 1).

**Fig. 2.**
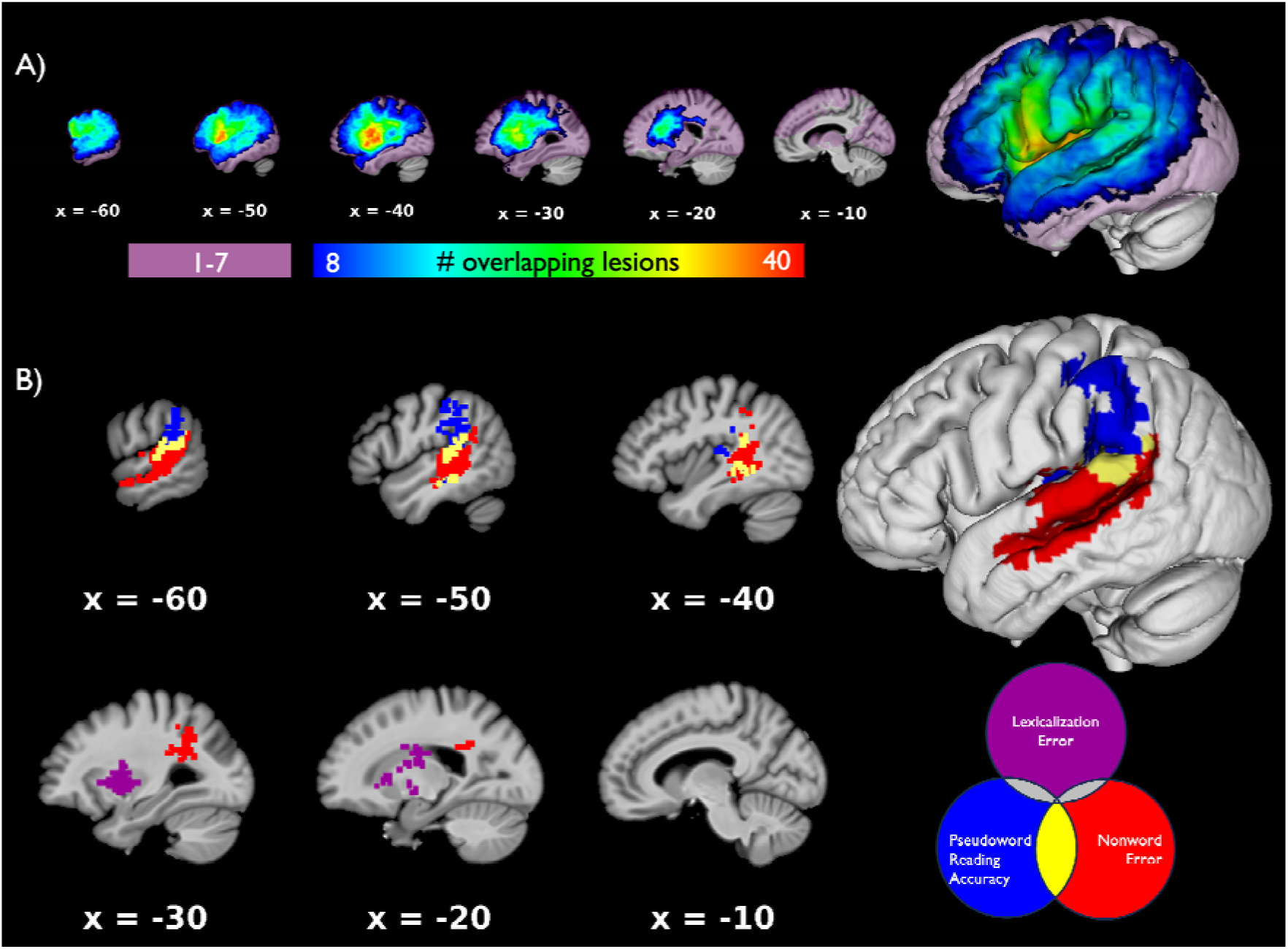
Lesion overlap map and SVR-LSM results at voxelwise *p*<0.005 and clusterwise *p*<0.05. **(A)** Light purple shading indicates voxels excluded from the SVR-VLSM analysis due to insufficient lesion coverage (<8). Color spectrum shading indicates voxels considered in the SVR-VLSM analysis. **(B)** SVR-VLSM analyses show that reduced pseudoword reading accuracy (blue) was associated with lesions to the posterior superior temporal gyrus and the supramarginal gyrus (clusterwise p=.009). Greater lexicalization error rate (purple) was associated with lesions to the left frontal white matter and anterior putamen (clusterwise *p*=0.043). Greater nonword error rate (red) was associated with lesions to the posterior superior temporal gyrus and sulcus (clusterwise *p*=0.002). Lesions associated with both nonword errors and pseudoword reading accuracy (yellow) were associated with the planum temporale and deep temporal white matter. There was no overlap between the lesions associated with lexicalization errors and either nonword errors or reading accuracy. Error rate analyses control for the alternate error rate, age, education, lesion volume, log-transformed chronicity in months, and Apraxia of Speech Rating Scale Apraxia Severity score.

**Table 4.**
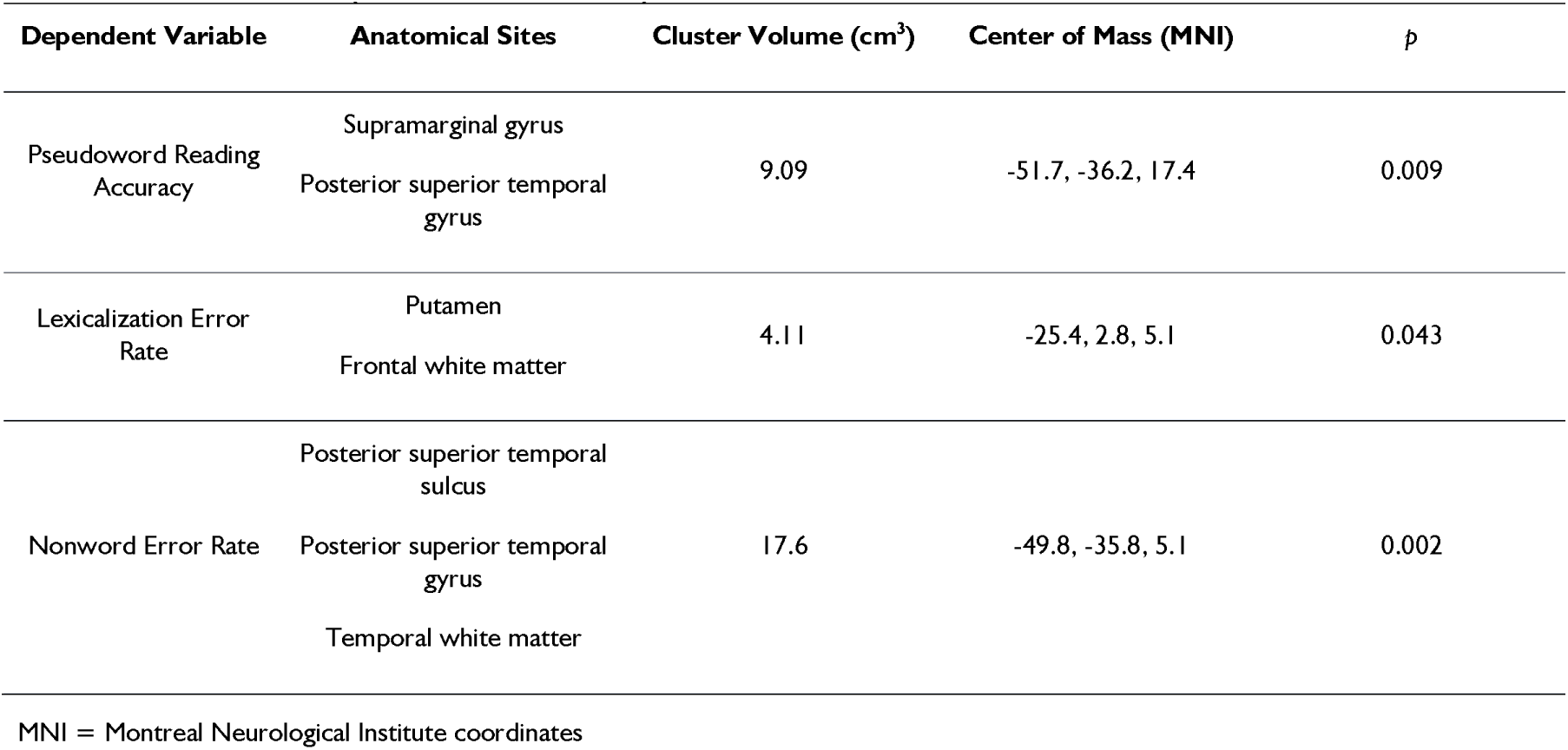
SVR-VLSM results (Cluster FWER P < 0.05)

| Dependent Variable | Anatomical Sites | Cluster Volume (cm<sup>3</sup>) | Center of Mass (MNI) | $p$ |
| --- | --- | --- | --- | --- |
| Pseudoword Reading Accuracy | Supramarginal gyrus | 9.09 | -51.7, -36.2, 17.4 | 0.009 |
|  | Posterior superior temporal gyrus |  |  |  |
| Lexicalization Error Rate | Putamen | 4.11 | -25.4, 2.8, 5.1 | 0.043 |
|  | Frontal white matter |  |  |  |
| Nonword Error Rate | Posterior superior temporal sulcus | 17.6 | -49.8, -35.8, 5.1 | 0.002 |
|  | Posterior superior temporal gyrus |  |  |  |
|  | Temporal white matter |  |  |  |

MNI = Montreal Neurological Institute coordinates

LE production was associated with lesions to the frontal white matter and left putamen (MNI centroid: −24.9, 1.1, 9.7; Fig. 2B; Table 4), which has been suggested as part of a subcortical pathway to send computed phonological codes to motor cortex.^70^ The unthresholded maps show that damage to the supramarginal gyrus, as well as motor and frontal cortex, were also weakly associated with increased LE rates (Supplementary Fig. 1).

In contrast, NE production was associated with a large VLSM cluster centered in the pSTS (MNI centroid: −49.8, −35.8, 5.1; Fig. 2B; Table 4). This cluster extended anterior along the STS, superior to the STG, and medially into the temporal white matter. The pSTS is likely involved in computing semantic contributions to oral reading.^40^ The unthresholded maps also show weaker association between NE error rate and damage to a broad temporoparietal network (Supplementary Fig. 1).

SVR-CLSM analysis identified 10 disconnections that impaired pseudoword reading accuracy (Fig. 3A, Supplementary Table 2). Seven disconnections were confined to the left SMG, STS, and STG. One disconnection between the SMG and insula, as well as one between the postcentral gyrus and superior temporal lobe, both suggest disconnections of the phonology-motor circuit. One disconnection of the middle occipital lobe and the insula suggests a role for visual processes.

**Fig. 3.**
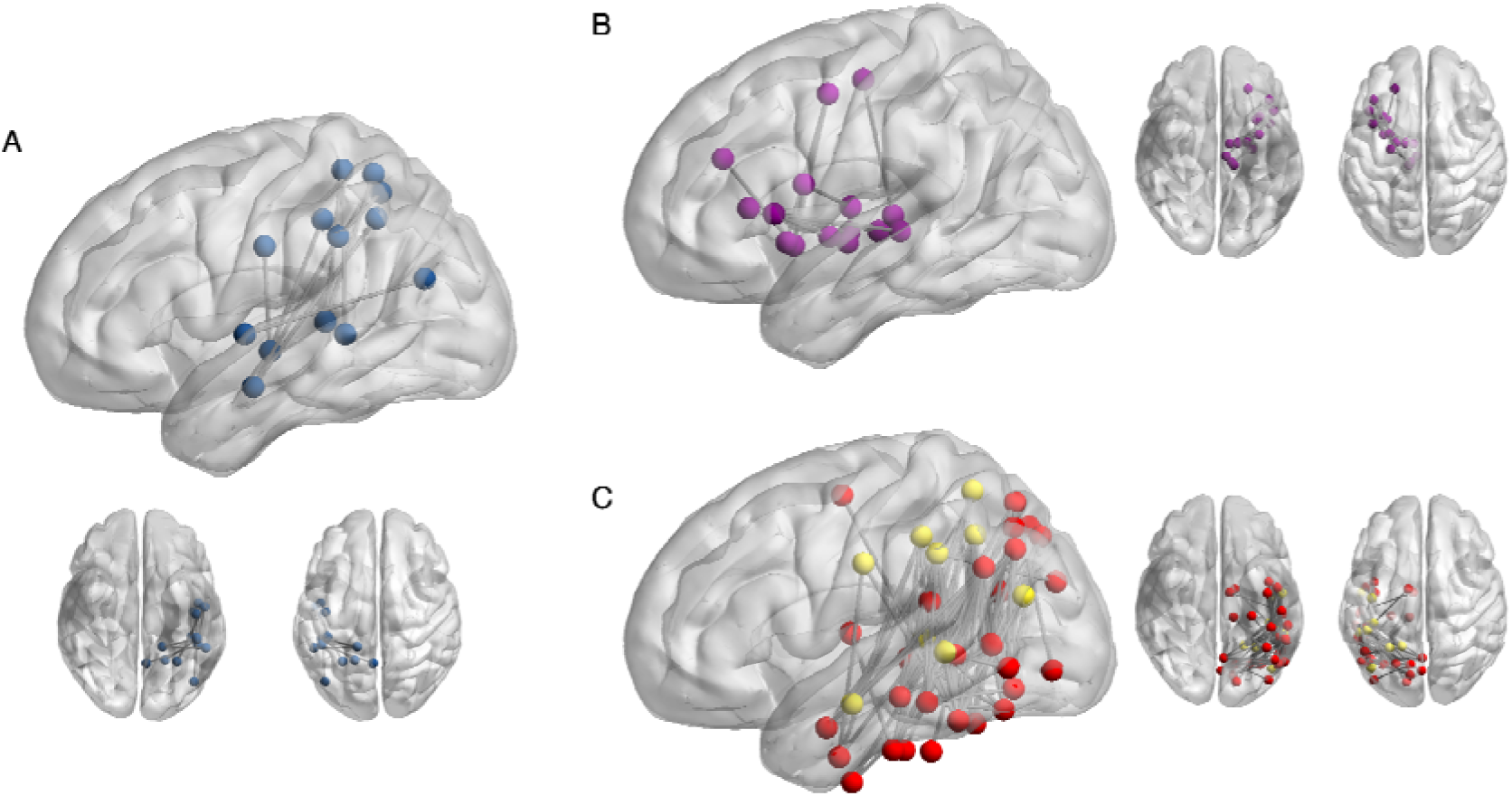
SVR-CLSM results at edgewise and parcelwise FWER < 0.05. **(A)** Disconnections that reduce pseudoword reading accuracy, controlling for age, education, and lesion volume. **(B)** Disconnections that reduce lexicalization errors. **(C)** Disconnections that reduce nonword errors. Nodes in yellow overlap with the pseudoword reading SVR-CLSM result. Nodes in red are unique to the nonword error analysis. Error rate analyses control for the alternate error rate, age, education, lesion volume, log-transformed chronicity in months, and Apraxia of Speech Rating Scale Apraxia Severity score. SVR-CLSM results visualized with BrainNet Viewer.^74^

SVR-CLSM analyses revealed that 84 disconnections, broadly temporoparietal, were associated with greater NE rates (Fig. 3B, Supplementary Table 3). Eight nodes, spanning the SMG, STS, and STG were involved in both the NE and pseudoword reading accuracy analysis. The remaining 29 nodes were broadly distributed in the temporal, parietal, and occipital cortex. Notably, disconnections of the anterior temporal lobe, the angular gyrus, and the fusiform/posterior inferior temporal gyrus are all cortical regions associated with lexical-semantic processing, supporting the behavioral findings.^71,72^ More posterior fusiform and occipital disconnections also indicate a potential role for visual and orthographic processing impairments in producing PE.

Increased rates of LE were related to 15 disconnections (Fig. 3C, Supplementary Table 4). Fourteen disconnections were between subcortical structures and frontal or motor cortex. The final disconnection was between the inferior frontal gyrus pars opercularis and a node in the middle frontal gyrus. These areas are strongly implicated in articulatory processing.^21,73^

### Identifying a shared neural substrate for lexical reading

Overlaying the SVR-VLSM map for NE errors with the results of Staples et al., 2025 demonstrated highly overlapping lesion maps, with lesions in the STS and STG producing both semantic reading deficits as evidenced by a reduced imageability effect and also high rates of NE in pseudoword reading (Fig. 4).

**Fig. 4.**
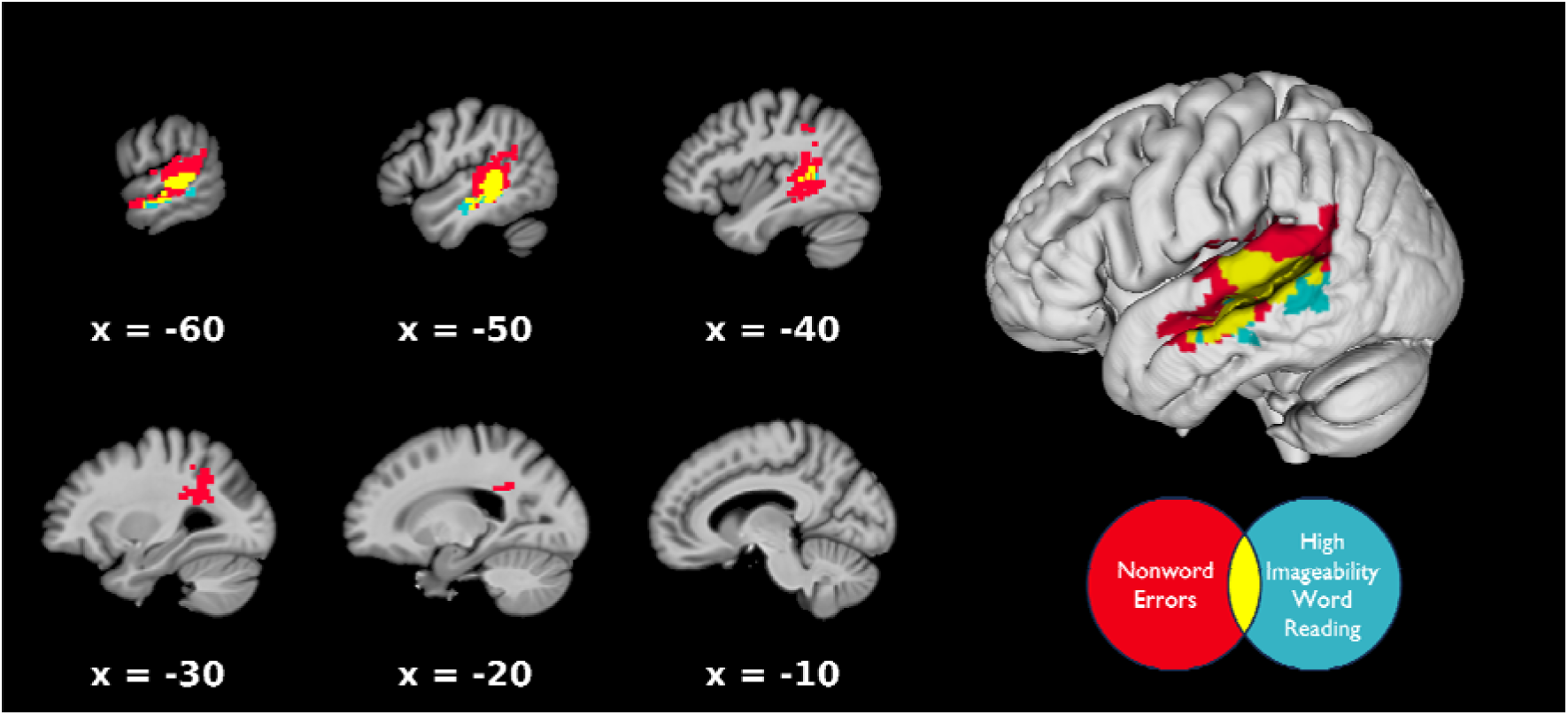
Superior temporal sulcus lesions reduce lexical-semantic contributions to reading. Overlap between the SVR-VLSM result for nonword error rate and the SVR-VLSM result that was found to reduce high imageability word reading, relative to low imageability word reading, in Staples et al.^40^

## Discussion

Leveraging a large sample of left-hemisphere stroke survivors, we identified the neurocognitive drivers of error production in pseudoword reading. Behaviorally, we found that pseudoword reading accuracy is predicted both by pseudoword repetition and naming ability and that most LHSS with alexia make both lexicalization and nonword errors, with great variability in relative error rates. Stroke survivors with more impaired pseudoword repetition produce more LE, while persons with more impaired naming make more NE. Greater rates of NE production are associated with lesions to the left superior temporal gyrus and sulcus and broad temporal, parietal, and occipital disconnection. LE are related to lesions to the left putamen and disconnections of the motor cortex. These results suggest 1) both lexical-semantic and phonological processes contribute to pseudoword reading in stroke alexia and 2) that error production in pseudoword reading reflects the relative intactness of lexical-semantic and phonological processing.

### Both phonological and semantic deficits contribute to pseudoword reading errors

Consistent with previous studies, LHSS in our sample were more phonologically than semantically impaired.^75^ Although pseudoword reading deficits are canonically interpreted as reflecting phonological reading deficits, i.e., phonological alexia, we found that both picture naming and pseudoword repetition impairments independently predicted pseudoword reading deficits, providing evidence for both lexical-semantic and phonological contributions. We found that most LHSS made both NE and LE, and that the rates of NE and LE errors were equivalent at the group level. Individual variation was large, however, and we found that the relative rates of NE and LE errors were subserved by different cognitive processes. Worse pseudoword repetition, but not picture naming, was associated with greater LE. This result stands in contrast with a previous finding that less impaired semantic processing, relative to controls, was correlated with higher rates of LE production.^41^ However, that study did not control for the contribution of phonological processing to lexicalizations. Assessing semantic deficits alongside phonological deficits is critical because pseudoword reading deficits are driven primarily by phonological impairment. Therefore, phonological deficits should lead to more errors overall, and the role of semantics in pseudoword reading must be in concert with phonology. When the semantic contributions are controlled for, we find that more impaired phonological processing increases LE.

In contrast, worse picture naming predicted greater NE rates when pseudoword repetition was controlled for. In an earlier paper, we found that the imageability effect in reading is related to the intactness of semantics-to-phonology mappings, as indexed by picture naming, rather than impaired semantic representation or control.^40^ As in that paper, we suggest that worse picture naming reflects an inability to use semantic information to influence phonology. Where stroke survivors with relatively intact naming can recruit lexical processes, and thus make LE, stroke survivors with impaired picture naming must rely on impaired sublexical assembly, as indexed by pseudoword repetition. The noisy output of this process increases NE production during pseudoword reading.

Consistent with this interpretation, we found that greater NE but not LE predicted worse high imageability word reading accuracy, relative to low imageability words. These findings directly link pseudoword reading and semantic contributions to real word reading, suggesting that damage to the lexical system systematically shifts reading towards a greater reliance on sublexical processing for all written letter strings. Unexpectedly, we also found that worse apraxia severity was associated with a larger imageability effect. While further targeted research is needed, this finding suggests that impaired motor-phonological processing leads to a greater reliance on semantic processes, when they are intact, and hence deficits in low imageability word reading with relative sparing of high imageability words.

### Identifying the neural loci of pseudoword reading errors

Lesions to the pSTG, STS, and temporal white matter increased NE rates. The STG is a critical dorsal stream processor. It is implicated in the representation of phonological information at every grain size, from the phonemic^14,15^ to the whole-word level^16,17^. The NE elicited in our study may result in part from a failure to access these representations, leading to downstream errors. The SVR-VLSM result for pseudoword accuracy does overlap with the result for NE in the planum temporale, suggesting that this area is a shared region for representing phonological units. Lesions along the STS were a unique neural correlate of NE production. The previous study by Staples et al. identified an overlapping region of the pSTS as a critical node for computing semantic influences in real word oral reading using an imageability manipualtion.^40^ Our results extend this previous finding: damage to the STS disrupts semantic contributions even during pseudoword reading.

Disconnections in a broad range of lateral temporal, parietal, and occipital cortex were also related to NE. A core set of disconnections in the SMG, STG, and MTG overlapped with disconnections that impaired pseudoword reading accuracy, replicating previous findings.^32^ The nodes that uniquely increased NE rates were broadly clustered in visual and ventral stream semantic cortex. The ATL and AG are both considered semantic ‘hubs’, although the role of each is debated.^76,77^ The middle FG has also been implicated as a multimodal naming region.^71^ Damage to these regions likely impairs core semantic representational access or processing. We also identify disconnections of MTG and ITG regions, inferior to the SVR-VLSM result, that have been previously associated with semantic control processes.^78,79^ Regardless of their exact roles, damage to semantic processors increases the rate of NE relative to LE. Finally, lesions to the ventral occipitotemporal (vOT) cortex are also uniquely related to NE. The vOT is a critical processor of orthographic information, suggesting that another possible source of NE errors is a failure correctly process the visual wordform.^80,81^

Increased LE rates were instead associated with lesions of the left putamen and surrounding white matter. Dynamic causal models suggest the putamen as part of an oral reading pathway, connecting the VWFA with the precentral cortex^70^. Dyslexic adults have reduced grey matter in the putamen, and lesions can cause aphasia.^82–84^ Meta-analytic connectivity modeling suggests that the left putamen may have a specific role in the semantic processing of visual words, and lesions to the putamen can cause semantic errors in oral reading and comprehension.^85,86^ Examination of the unthresholded SVR-VLSM maps additionally show a sub-threshold peak in the SMG, reinforcing the role of sublexical processing in increased error rates in pseudoword reading. The unthresholded maps also show weaker associations with increased LE rate throughout the basal ganglia and the posterior IFG. SVR-CLSM of LE rates identified a network of disconnections between the frontal cortex and the insula, putamen, thalamus, and globus pallidus that increased LE rate. These subcortical structures are critical to both speech planning and motor loops.^21,73^ Together, these results suggest that damage to dorsal stream regions involved in sublexical assembly for speech production leads to a greater reliance on the well-learned motor plans associated with real words, and thus to LE.

### A deficit-based model of phonological and semantic influences on pseudoword reading

Taken together, these data suggest that the reading of all pronounceable letter strings, including pseudowords, is influenced both by phonological and semantic processes, at least in post-stroke alexia. These results are consistent with a model in which the extent of damage to one stream (e.g., the ventral stream) impacts the degree to which the other (the dorsal stream) is relied upon when reading pseudowords. Our data cannot definitively adjudicate between an account where these influences are due to an immediate switch to reliance on the less-damaged processor or one where plasticity-based recovery strengthens the less-damaged processor. However, chronicity was a significant predictor of LE production and a marginally significant predictor of NE production. This result points to recovery-related reorganization of remaining neurocomputational resources as a potential mechanism. Longitudinal studies tracking error production from the acute stage into the chronic stage after stroke would help to confirm this interpretation.

This process of dynamic, plasticity-related recovery is predicted by artificial neural network models of alexia recovery. Immediately after damage to these models, performance is globally poor due to diaschisis-like network dysfunction.^43^ As the models are trained to simulate recovery, the division of labor between the orthography-to-phonology pathway and the semantically-mediated pathway is altered. Phonological damage causes the intact semantic pathway to become more involved in processing inputs, producing lexicalizations.^43^ Other computational studies have examined regularization errors, which occur when semantic processes are impaired and the network comes to rely on the orthography-to-phonology pathway.^9,34^ While the impact of semantic damage and recovery on pseudoword reading has not been tested in these models, our data predicts that the ratio of NE to LE would increase. However, we note that these models do not explicitly accommodate motor processing. Integrating computational models of reading with extant models of speech production^87^ may be essential to accurately modeling the effects of lesions and plastic network changes on pseudoword reading.

### Clinical implications

Our findings have important implications for how pseudoword reading performance is interpreted when diagnosing alexic deficits in order to plan for treatment. The findings suggest that semantic deficits may play a role in poor pseudoword reading, such that targeting semantics during alexia treatment might improve pseudoword reading. The findings may also aid in interpreting errors made during behavioral interventions for alexia, particularly in approaches such as phonomotor therapy which rely on retraining articulatory precision and phonological assembly skills.^88^ Persons with greater phonological impairments may benefit from attempting to strategically avoid producing real words when they are training grapheme-phoneme combinations. Finally, these results may have implications for preoperative language mapping for resections due to tumors, epilepsy, or other debilitating neurological disorders. In line with the focus of neurocognitive research on aphasia rather than alexia, picture naming is the most widely used task for identifying eloquent cortex.^89^ Because pseudoword reading is likely to induce errors, and because we show that these errors can be mapped onto underlying cognitive function, including a pseudoword reading task during presurgical language mapping may be a time-efficient and effective way to prevent the removal of tissue that would cause subtle language disorders.

### Limitations

Our results have important limitations. Lesion distribution in our sample was typical of MCA stroke, and thus did not include sufficient coverage of the left ventral occipitotemporal cortex (lvOT), a critical node for processing orthographic forms, for SVR-VLSM.^81,90^ However, we did find that disconnections along the length of the vOT caused NE. In addition, a fundamental limitation is that LSM techniques identify only cortex or disconnections that are very strongly associated with a behavior, and so cortical regions with less strong associations or that are involved in widely distributed functions are less likely to be identified. For this reason, we presented unthresholded SVR-VLSM maps in addition to the statistically significant results, but the patterns in these unthresholded maps must be interpreted cautiously. Finally, the approach used here is associational and individual stroke survivors may not adhere to the lesion or behavioral patterns described here.

## Conclusions

We provide evidence that pseudoword reading deficits relate not only to phonological deficits, but to deficits in mapping semantics to phonology as well. The types of errors produced in pseudoword reading are influenced by damage to semantic and phonological processes and the brain regions that subserve them. These data argue for ventral stream semantic modulation of dorsal stream phonological processing in pseudoword reading, constraining neurocognitive theories of reading and alexia recovery.

## Data availability

Data may be made available upon request.

## Acknowledgements

We thank the following individuals who contributed to data collection, in alphabetical order: Elizabeth Dvorak, Trini Kelly, Elizabeth Lacey, Sachi Paul, and Candace van der Stelt.

## Funding

This work was supported by National Institute on Deafness and Other Communication Disorders: R01DC014960 and R01DC020446 to PET, R00DC018828 to ATD, and T32DC019481 to RS.

## Competing interests

The authors report no competing interests.

## Notes

### Competing Interest Statement

The authors have declared no competing interest.

### Summary of Updates

Added author after helpful comments; author affiliations updated

