## Supplementary Materials for "Meaning for reading pseudowords: errors reveal semantic influences on pseudoword reading after stroke"

Supplementary Material

Supplementary Table 1 – Results of a linear mixed effects model pseudoword repetition and naming in stroke survivors.

| Variable | df | SS | MSE | F | *p* |
| --- | --- | --- | --- | --- | --- |
| **Task** | **1** | **0.12** | **0.08** | **4.71** | **.033** |
| Age | 1 | 0.09 | 0.08 | 1.12 | .291 |
| Education | 1 | 0.02 | 0.08 | 0.20 | .655 |
| **Lesion Volume (cm^3^)** | **1** | **1.21** | **0.08** | **15.58** | **<.001** |
| **Chronicity in months (log-transformed)** | **1** | **0.44** | **0.08** | **5.84** | **.018** |
| **ASRS3 Apraxia of Speech Severity** | **1** | **1.42** | **0.08** | **18.58** | **<.001** |
| Age:Task | 1 | 0.04 | 0.03 | 1.71 | .195 |
| Education:Task | 1 | 0.02 | 0.03 | 1.03 | .314 |
| Chronicity:Task | 1 | 0.03 | 0.03 | 1.18 | .280 |
| ASRS3:Task | 1 | 0.01 | 0.03 | 0.34 | .561 |
| Lesion Volume:Task | 1 | 0.04 | 0.03 | 1.69 | .197 |

Supplementary Table 2 – SVR-CLSM results identifying disconnections associated with reduced pseudoword reading accuracy.

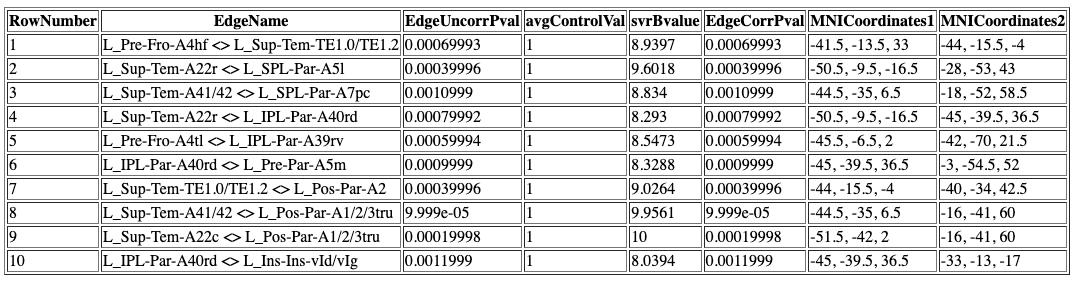

Supplementary Table 3 – SVR-CLSM results identifying disconnections associated with reduced pseudoword reading accuracy.

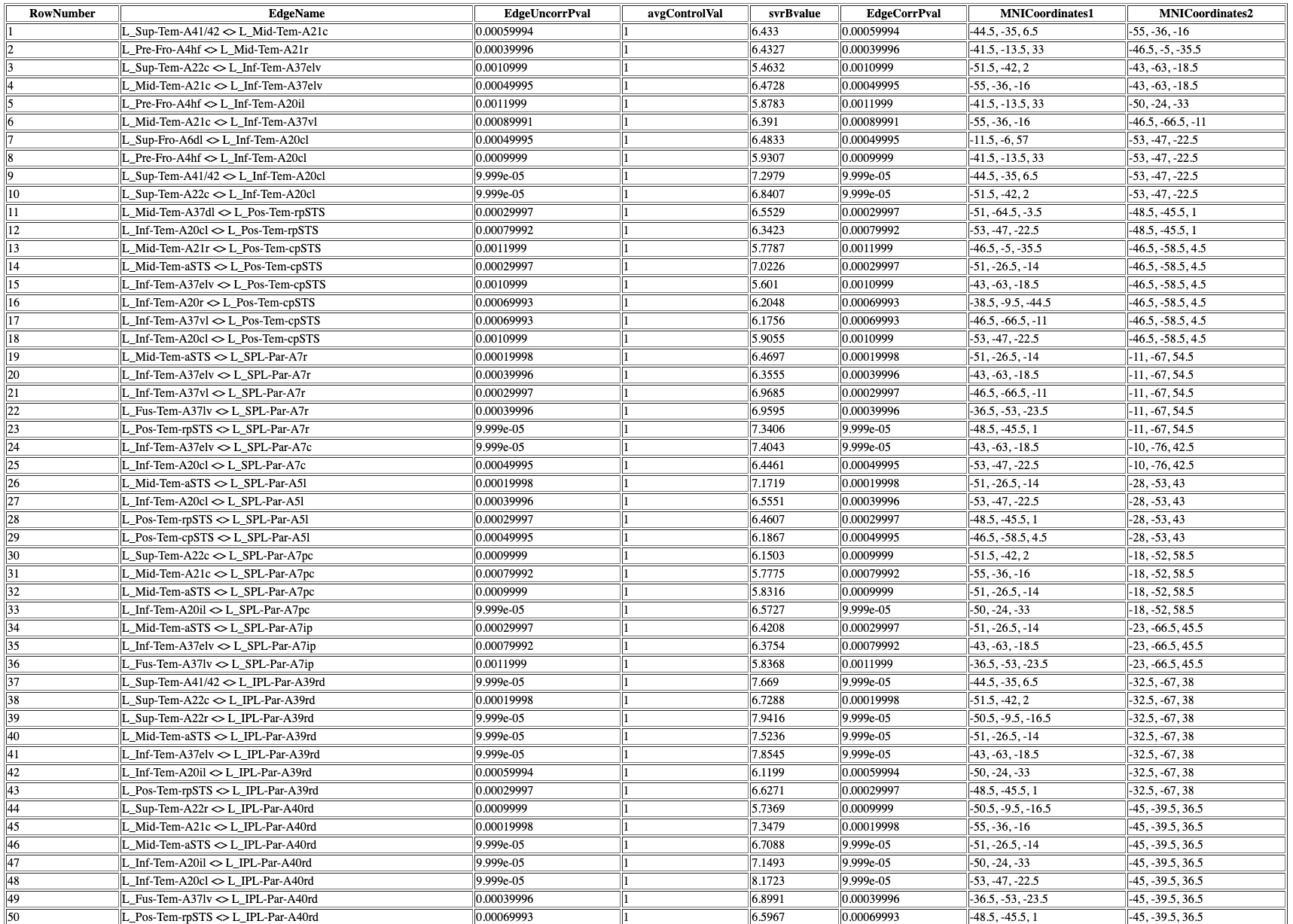

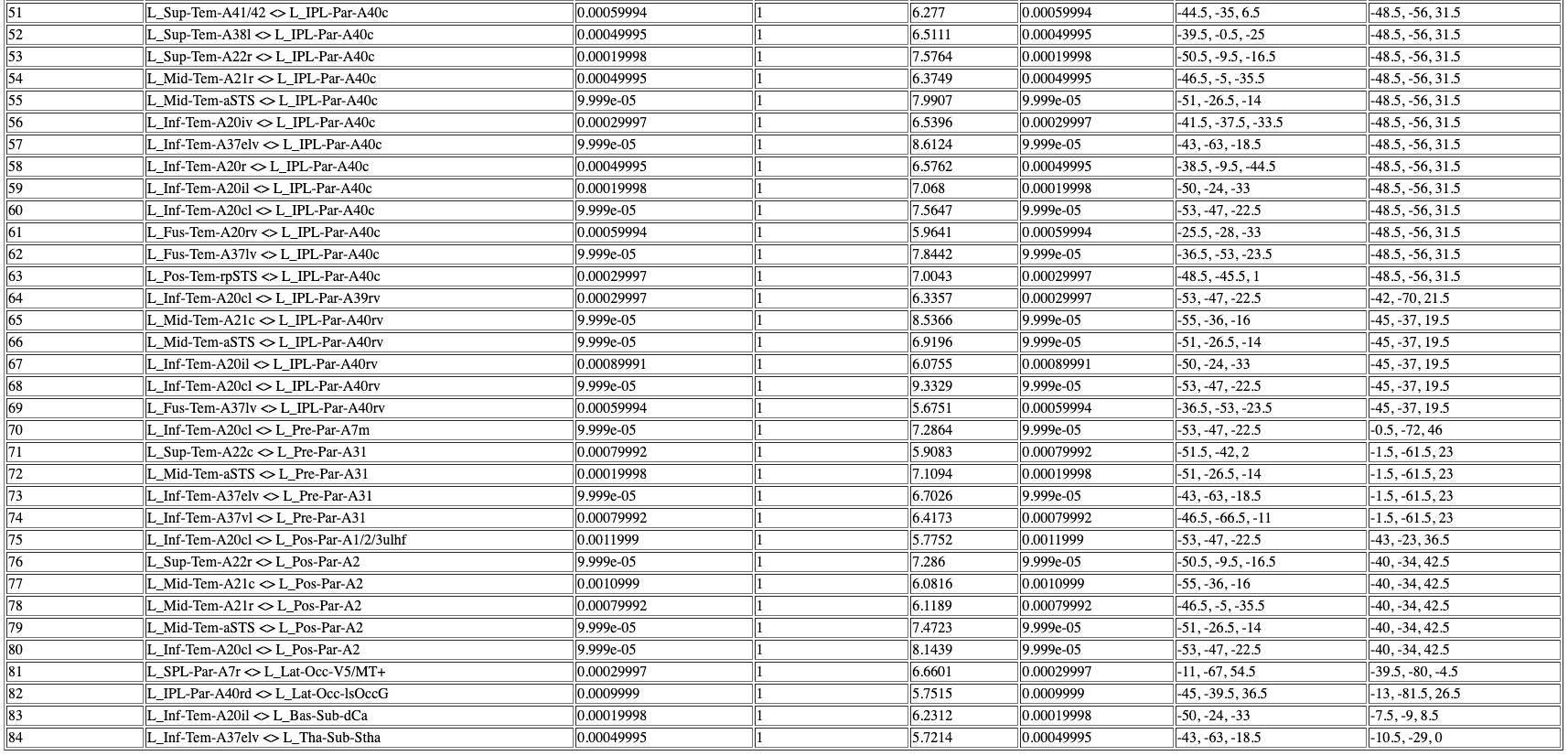

Supplementary Table 4 – SVR-CLSM results identifying disconnections associated with reduced pseudoword reading accuracy.

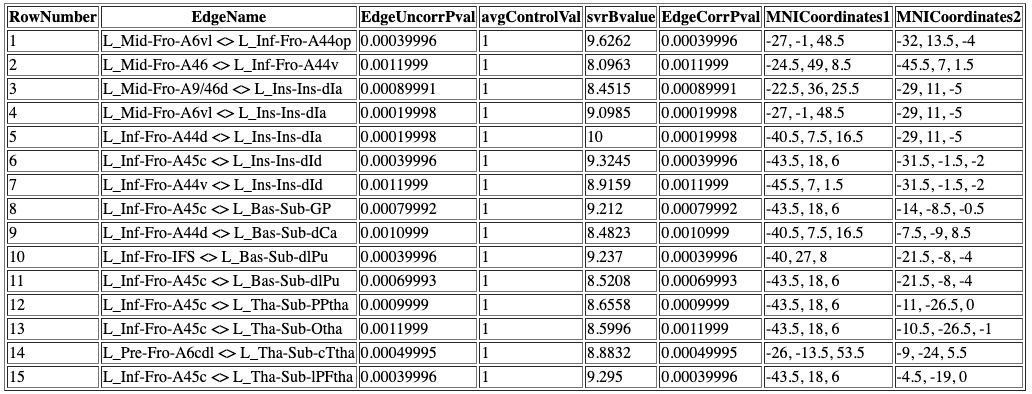

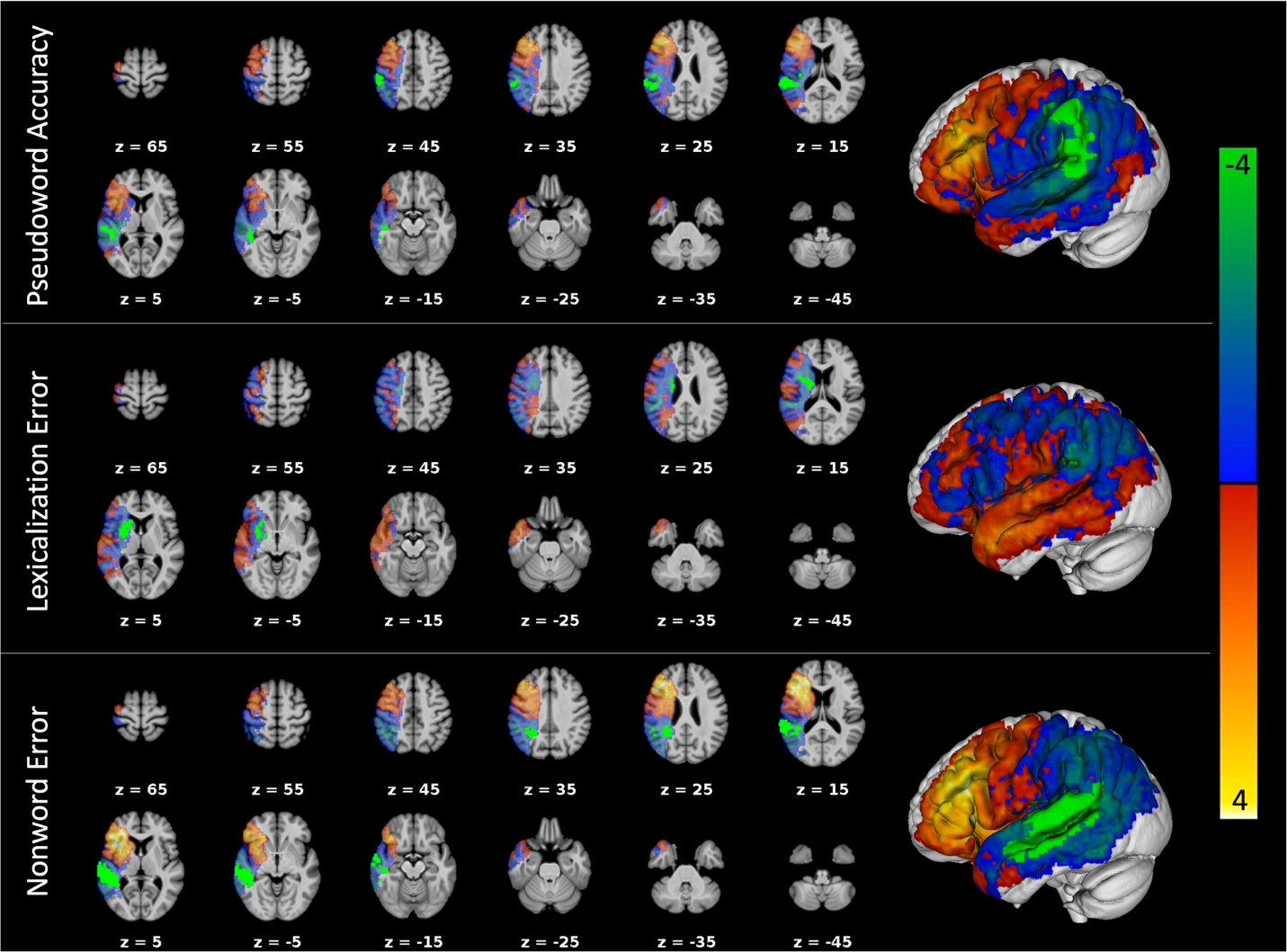

Supplementary Fig. 1 – Unthresholded Z score maps for the three VLSM analyses, demonstrating the subthreshold patterns associated with each score. Opaque light green indicates the significant SVR-VLSM finding. Transparent green-yellow spectrum indicates Z-scores.
